# Behavioural and physiological evidence for the development of cardiac-exteroceptive integration during the first year of life

**DOI:** 10.1101/2025.07.31.668000

**Authors:** Tomoko Isomura, Ayami Suga, Megumi Kobayashi, Yuri Terasawa, Kenta Kimura, Hideki Ohira

## Abstract

Continuous integration of environmental exteroceptive and internal interoceptive signals is fundamental for perception and adaptive behaviours, yet its developmental trajectory remains poorly understood. Here, we introduced a modified ‘iBEATs’ paradigm, based on prior work, to investigate cardiac–exteroceptive integration in 3–8-month-old infants. Using behavioural measures of looking time and physiological measures of pupillometry, we found that the ability to detect cardiac–exteroceptive synchrony emerges during the first year of life. Critically, this was evident only when stimuli coincided with **s**ystole, the baroreceptor-active phase of the cardiac cycle, supporting central multimodal integration interpretable within a predictive coding framework. Furthermore, individual differences in behavioural sensitivity were accounted for by the degree of autonomic maturation. These findings provide the first evidence for the developmental emergence and mechanisms of cardiac-exteroceptive integration, and establish the modified iBEATs paradigm as a promising tool for assessing interoceptive development in early life.

---

Maintaining physiological states within specific ranges (i.e. homeostasis), and adaptively regulating them (i.e. allostasis), are essential for the survival of organisms, including humans, who have, therefore evolved the capacity to monitor and regulate their internal bodily states. The central processing of signals originating from within the body is referred to as interoception. Given its close ties to vital functions, interoception is thought to serve as a driving force in shaping perceptions of, and actions toward, the external world. Crucially, the dynamic integration of interoceptive signals with exteroceptive cues—information derived from the environment—is fundamental for evaluating sensory inputs and guiding adaptive behaviours.

Experimental research has predominantly focused on the temporal coupling between interoceptive signals and external stimuli. For instance, the heartbeat discrimination task—participants judge the synchrony between heartbeats and external stimuli, such as sounds or lights—has been widely used for assessing the ability to perceive interoceptive signals (Hickman et al., 2020; Whitehead et al., 1977). In this task, synchronous stimuli are typically presented 200–300 ms after the R-wave of an electrocardiogram (ECG), a delay chosen to coincide with the cardiac *systole*. Following ventricular depolarization—represented by the R-wave—the ventricles contract and expel blood from the heart. Baroreceptors located in the aortic arch and carotid sinuses detect this surge in blood pressure, and convey information about the timing and intensity of the heartbeat to the central nervous system via the vagal and glossopharyngeal nerves (Brown & Eccles, 1934; Levy & Martin, 1984). The systolic period, occurring 200–300 ms, after the R-wave, thus, corresponds to the central representations of cardiac afferents’ peak—a moment when heartbeat sensations are most accessible to awareness, and exert the strongest influence on perceptual and cognitive processes (Brener & Kluvitse, 1988; Pollatos & Schandry, 2004; Yates et al., 1985). Numerous studies have demonstrated that a range of perceptions and cognitions are modulated by the timing of external stimuli relative to the cardiac cycle (Garfinkel & Critchley, 2016; Skora et al., 2022).

More recently, researchers have begun exploring the developmental origins of interoceptive processing. To probe infants’ sensitivity to heartbeat-exteroceptive synchrony, a pioneering study by Maister et al. (2017), introduced a task named ‘iBEATs’—comprising an animated character that moves either in sync (synchronous) or out of sync (asynchronous) with infants’ heartbeat—and compared infants’ looking times for each. They found that five-month-olds looked significantly longer at asynchronous stimuli, suggesting that they could detect heartbeat-exteroceptive synchrony. However, subsequent studies—although their procedures varied slightly—have produced mixed findings. Some replicated the original effect (Charbonneau et al., 2022 with rhesus macaques; Imafuku et al., 2023), whereas others failed to do so (Tünte et al., 2025; Weijs et al., 2023; see Donaghy et al., 2025 for a review). These divergent outcomes question the robustness of iBEATs in its original form. One potential issue involved in the original iBEATs task is the timing of the stimulus presentation. In all previous iBEATs studies, the ‘synchronous’ stimuli were time-locked to the ECG R-wave—a moment of minimal baroreceptor activity—which likely renders heartbeat–exteroceptive integration suboptimal, and thus undermines its accurate assessment. Furthermore, previous studies provide limited information beyond mean looking times, and their participants’ ages varied (e.g. 5 months in Maister et al., 2017; 6 months in Imafuku et al., 2023; 5–7 months in Weijs et al., 2023; and 3, 9, and 18 months in Tünte et al., 2025), making it difficult to interpret the findings from a coherent developmental perspective (see Donaghy et al., 2025, for a review). Addressing these methodological issues, and consolidating the experimental procedures, is crucial for advancing research on interoceptive development.

We introduced a refined iBEATs paradigm (Fig. 1) to examine how infants integrate cardiac and exteroceptive signals, and how this integration shapes their behaviour. As our key manipulation concerns stimulus timing, we implemented two synchrony conditions by locking the stimulus either to: 1) mid-systole (200–300 ms after the R-wave, when the central representation of cardiac signals peaks), or 2) diastole (at the R-wave itself, when baroreceptor input is minimal). Contrasting each synchrony condition with an asynchronous control allowed us to isolate the contribution of the central cardiac representation from a mere rhythmic match, yielding a more sensitive measure of infants’ cardio-visual integration. Furthermore, to illuminate the mechanisms underlying any looking-time differences, we recorded task-evoked pupil dilation while infants viewed the animations, as pupil size is thought to index information gain (Zénon, 2019), and been shown to increase during multimodal integration (Rigato et al., 2016; Wang et al., 2017). Further, interpreting these pupil responses within a predictive-coding framework—as a precision-weighted prediction-error signal—allows linking infants’ looking behaviour to the computational principles of predictive inference, which are now widely recognised in cognitive neuroscience. We focused on infants aged 3–8 months to track developmental changes during their first year of life. Converging evidence suggests that approximately 6 months marks a critical period of transition in neural system development, and is thought to support cardiac-exteroceptive integration (Atzil et al., 2018; Gao et al., 2015, 2009; Izard et al., 1991; Porges & Furman, 2011). In particular, the default-mode network, salience network, and vagally mediated autonomic pathways are largely immature at birth, but undergo rapid structural and functional maturation by roughly 6 months of age (Atzil et al., 2018; Gao, Alcauter, Smith, et al., 2015; Gao et al., 2009; Izard et al., 1991; Porges & Furman, 2011). By sampling across this window, we aimed to capture the emergence and early consolidation of cardio-visual integration. To test whether autonomic maturation contributes to this process, we also assessed the vagal function using infants’ resting heart rate variability (HRV).

**Figure 1.**
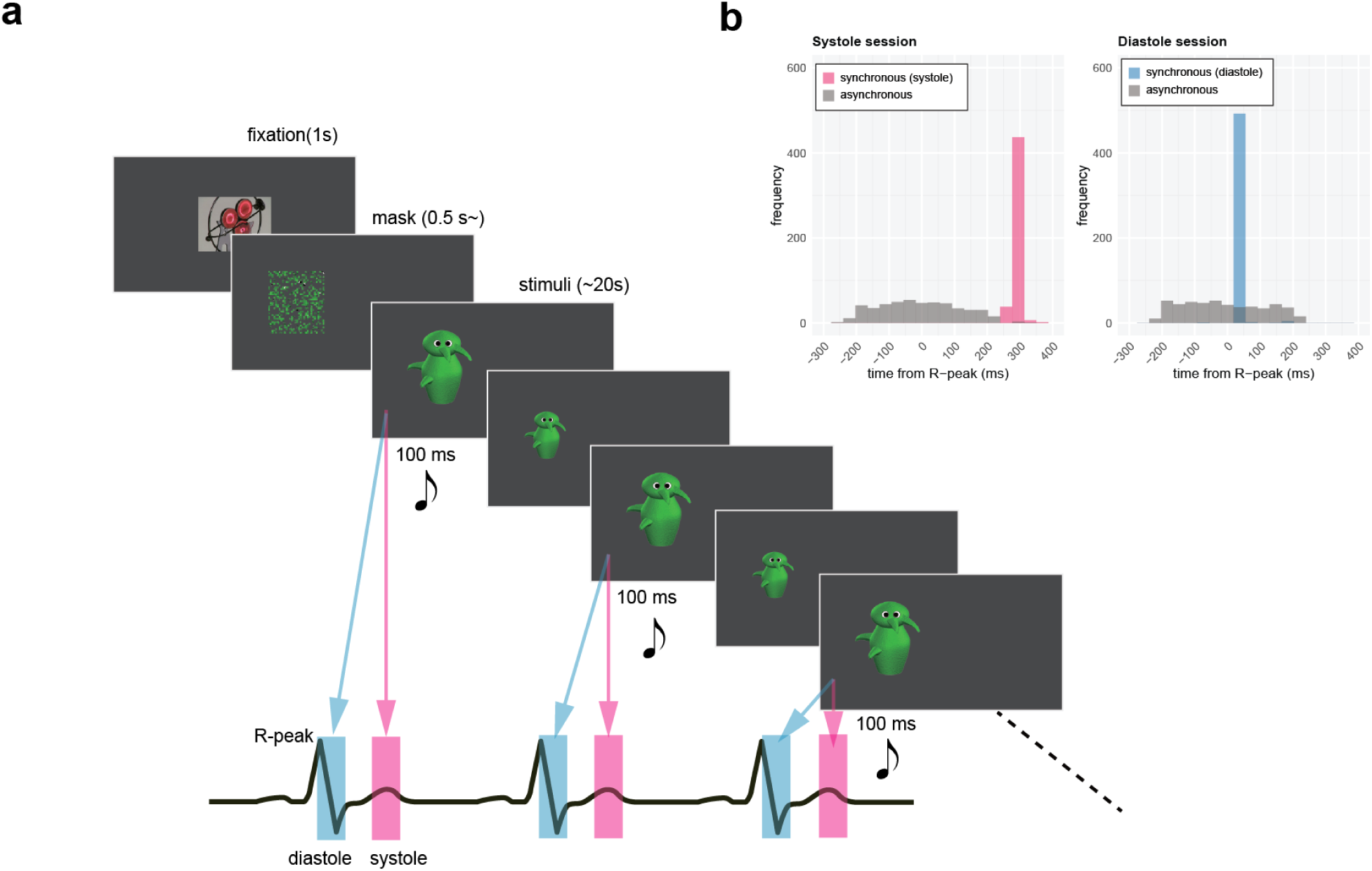
Modified iBEATs paradigm and manipulation of stimulus timing. **a.** Schematic of a synchronous trial in the modified iBEATs paradigm. Infants viewed an animated character—whose appearance varied across trials—moving in sync with their own heartbeat by briefly enlarging for 100 ms at a target cardiac phase (systole or diastole in separate sessions), thereafter shrinking back until the next enlargement. Each movement was accompanied by a fixed auditory tone. Trials ended when the infant looked away for more than 3 s or after 20 s. Synchronous and asynchronous trials—the latter using pulses drawn from other infants and adjusted by ± 10 % to match the infant’s own preceding rate—were presented in strict alternation, and stimulus position (left/right) were also alternated; both trial type and position were fully counterbalanced across infants. **b.** Example of a participant’s histogram having actual stimulus-onset times relative to the ECG R-wave. In the systole session (left), synchronous stimuli were targeted to be presented 250–300 ms after the R-peak (pink); in the diastole session (right), they were targeted to be presented 0–50 ms after the R-peak (blue). Asynchronous trials drew timings from other infants’ heartbeat recordings, producing an approximately uniform distribution across the cardiac cycle (grey).

## Results

Thirty-six infants (3–8 months/100–262 days-old; 23 girls) completed the modified iBEATs paradigm (Fig. 1A). Each infant was tested twice on separate days—once in a systole session and once in a diastole session. During both sessions, they viewed an animated character that pulsed (expanded) either *synchronously* with their own heartbeat, or *asynchronously*. The experiment thus followed a 2 × 2 design that crossed the cardiac phase (systole versus diastole) with temporal contingency (synchronous versus asynchronous). In synchronous trials, the pulse was phase-locked to mid-systole (R + 250–300 ms) in the *systole* session, and near the R-wave (R + 0–50 ms) in the *diastole* session (Fig. 1B). In asynchronous trials the pulse sequence was drawn from recordings of other infants (data collected in Suga et al., 2019) and rescaled by ± 10 % relative to the infant’s own inter-beat interval from the preceding trial, thereby eliminating real-time contingency. To minimise the influence of low-level preference, the character’s shape and colour varied randomly across trials, and each flash was accompanied by a constant auditory tone—an additional refinement over earlier iBEATs implementations. Synchronous and asynchronous trials were alternated, and characters appeared alternately on the screen’s left or right. A trial ended if the infant looked away for more than 3 s, or after 20 s. Each session continued for a maximum of 60 trials, or until the infant became too tired or fussy to continue. Throughout both sessions, infants’ gaze locations and pupil diameters were recorded. Finally, a 5-minute resting-state ECG was collected while infants lay quietly, providing an index of vagally mediated HRV for each participant.

### Looking behaviour

We first examined the infants’ looking behaviour. Across the two sessions, the infants completed a variable number of trials (systole session: *M* = 42.5, Standard Deviation [*SD]* = 13.5, range = 18–59; diastole session: *M* = 36.7, *SD* = 13.4, range = 18–59). As expected, looking times declined over the trials, indicating habituation (Fig. 2A). To illustrate developmental differences, data were plotted separately for older infants (≥ 6 months, 182–262 days old; n = 18) and younger infants (< 6 months, 100–180 days-old; n = 18).

**Figure 2.**
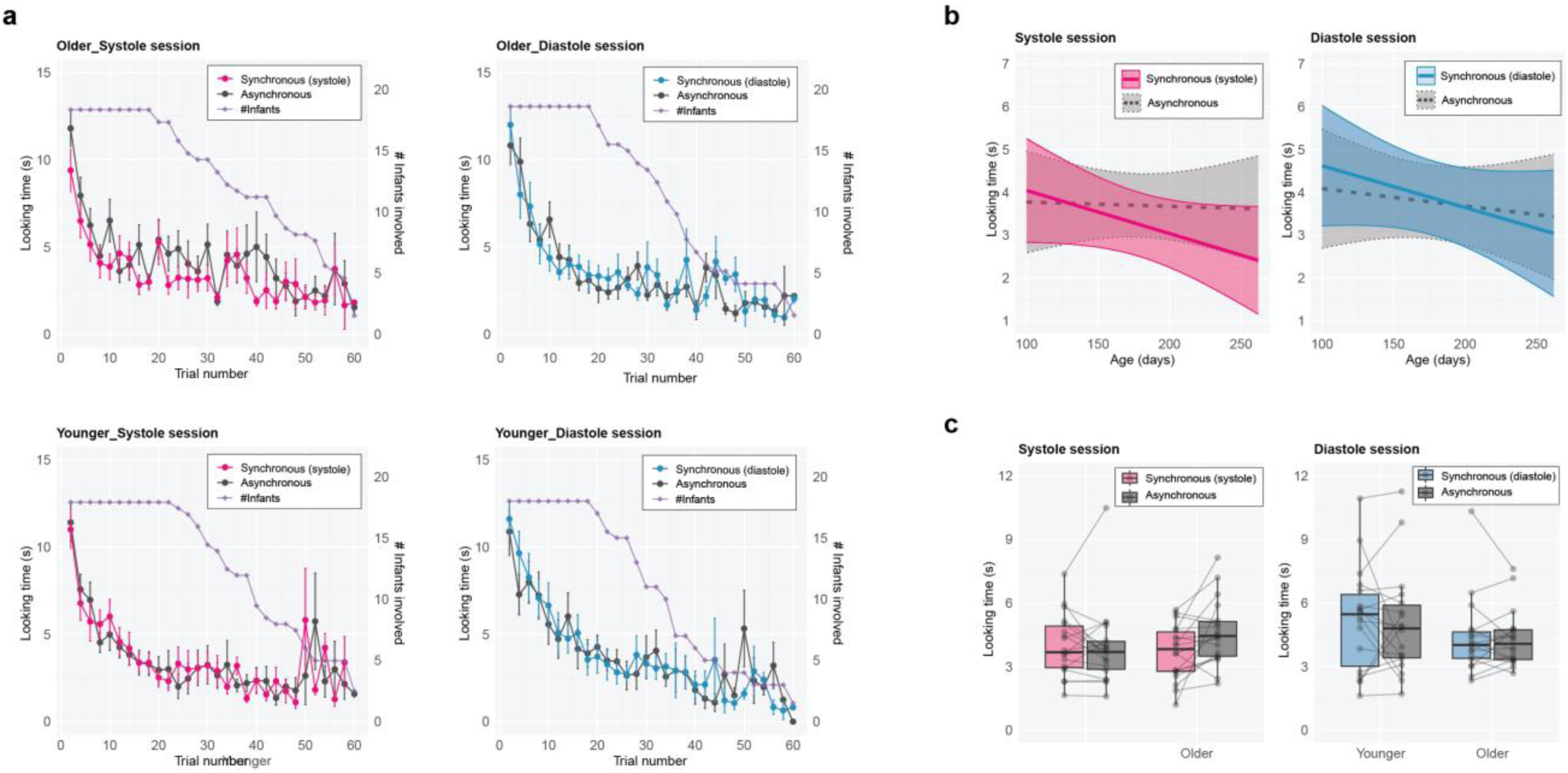
Results of the looking time. **a.** Mean looking time across trial numbers, for the systole session (left) and diastole session (right), plotted separately for older (≥ 6 months; top) and younger (< 6 months; bottom) infants (n = 18 each). As the synchronous and asynchronous trials alternated, within each trial-pair (1-2, 3-4, …), one trial was synchronous and the other asynchronous; looking times for each condition are plotted at the corresponding trial-pair index. Pink shows systole-locked synchronous trials, blue shows diastole-locked synchronous trials, and grey shows asynchronous trials. The purple line (right y-axis) indicates the number of infants contributing data at each trial-pair, reflecting attrition as infants became tired or fussy. **b.** Model-predicted looking times as a function of age from linear mixed-effects models, shown separately for the systole session (left) and the diastole session (right). Solid lines denote synchronous trials (pink for systole-locked, blue for diastole-locked), and dashed grey lines denote asynchronous trials. Shaded bands represent 95 % confidence intervals. **c.** Mean looking time averaged across all trials, shown separately for the systole session (left) and diastole session (right). Boxplots depict synchronous (pink for systole, blue for diastole) and asynchronous (grey) conditions; individual data points are overlaid.

To test the effects of stimulus synchrony and age, we fitted a linear mixed-effects model (LMM) to the systole-session data, with looking time as the outcome, and fixed effects of synchrony (synchronous versus asynchronous), z-scored age, and their interaction, plus random intercepts for participant and trial number (looking time ∼ synchrony + age + synchrony:age + (1|participant) + (1|trial)). This full model outperformed a null model with only random effects (*χ2*(3) = 12.60, *p* = 0.0056), and a model with the main fixed effects only (*χ2*(1) = 5.17, *p* = 0.023). Crucially, there was a significant synchrony × age interaction (*β* = –0.39, *t*(1358.4) = –2.27, *p* = 0.023) alongside a main effect of synchrony (*β* = –0.44, *t*(1365.3) = –2.56, *p* = 0.011; Table 1). A follow-up contrast confirmed that the age-related decline in looking time was steeper for synchronous than for asynchronous stimuli (*t*(1360) = 2.27, *p* = 0.023), indicating that older infants looked proportionally less at synchronous stimuli (Fig. 2B, left). Separate LMMs within each age-group showed the synchrony effect only in older infants (β = –0.94, *t*(663.6) = –3.69, *p* < .001) and not in younger infants (β = 0.11, *t*(655.7) = 0.49, *p* = 0.62). The raw distributions of individual mean looking times (Fig. 2C, left) mirror the mixed-model results: older infants looked longer in asynchronous than in synchronous trials, whereas younger infants showed no difference.

**Table 1.** Results of the linear mixed model predicting looking time.

| | $\beta$ | s.e. | t-value | df | p-value |
| --- | --- | --- | --- | --- | --- |
| *Systole condition* |  |  |  |  |  |
| intercept | 3.69 | 0.38 | 9.8 | 83.8 | < 0.001 |
| **synchrony (sync vs async)** | **-0.44** | **0.17** | **-2.6** | **1365.3** | **0.011** |
| age | -0.043 | 0.26 | -0.16 | 41.6 | 0.87 |
| **synchrony × age** | **-0.39** | **0.17** | **-2.3** | **1358.4** | **0.023** |
| *Diatole condition* |  |  |  |  |  |
| intercept | 3.78 | 0.44 | 8.6 | 84.8 | < 0.001 |
| synchrony (sync vs async) | 0.10 | 0.19 | 0.54 | 1167.0 | 0.59 |
| age | -0.18 | 0.31 | -0.57 | 39.6 | 0.58 |
| synchrony × age | -0.25 | 0.19 | -1.3 | 1165.6 | 0.19 |

In the diastole session, an analogous LMM did not improve fit over simpler models (full versus null: *χ2*(3) = 3.04, *p* = 0.39; full versus main-effects-only: *χ2*(1) = 1.76, *p* = 0.18), and neither synchrony nor age predicted looking time (Table 1; Fig. 2B–C, right).

Together, these results indicate that sensitivity to cardiac–exteroceptive synchrony emerges during the first year of life, but only when stimuli coincide with the baroreceptor-active window of systole—supporting a role for central representation of cardiac signals rather than mere rhythmic matching. More detailed data (e.g. fast versus slow asynchronous rhythms) are presented in Table S1.

### Pupil Responses

Next, we analysed infants’ pupillary responses to probe the mechanisms behind their looking behaviour. Given that, pupils could only be measured when infants fixated on the screen, and fixation declined over time within and across trials, we restricted our analysis to the first 4 s after stimulus onset in the first 12 trials (6 synchronous, 6 asynchronous), a window in which > 70 % of pupil samples were successfully tracked. The pupil size was baseline-corrected in each trial using the mean diameter during the 200 ms (mask presentation) immediately preceding stimulus onset. Fig. 3A shows the mean change in pupil size within this window, plotted separately for older and younger infants.

**Figure 3.**
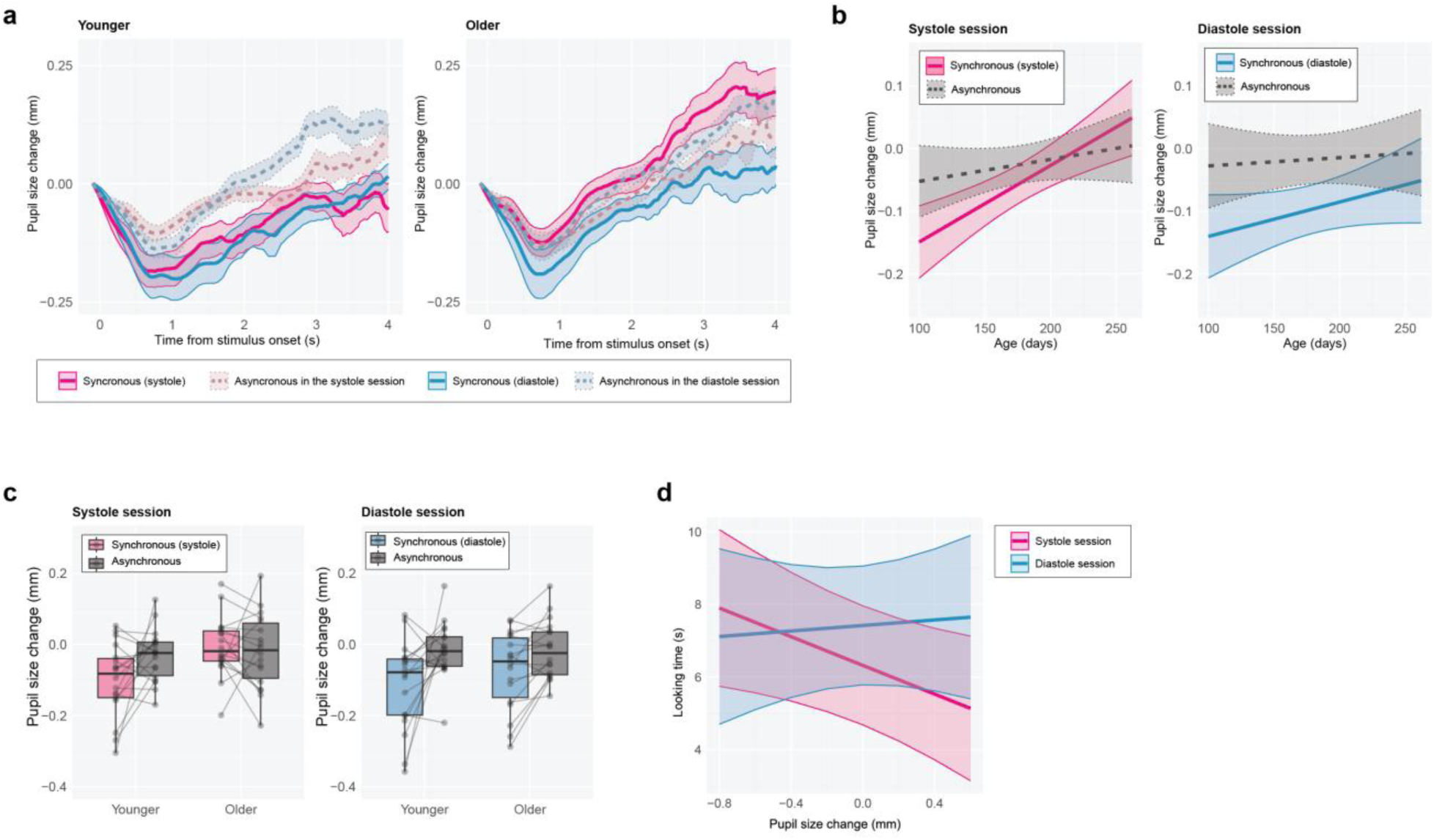
Results of pupil responses. **a.** Grand mean time course of baseline-corrected pupil size over the first 4 s following stimulus onset, shown separately for younger (left) and older (right) infants. Solid lines denote synchronous trials (pink for systole-locked; blue for diastole-locked), and dashed lines denote asynchronous trials in the corresponding session (pink for systole; blue for diastole). Shaded bands indicate ± 1 SEM. **b.** Model-predicted pupil dilation as a function of age from linear mixed-effects models, shown separately for the systole session (left) and the diastole session (right). Solid lines denote synchronous trials (pink for systole-locked, blue for diastole-locked), and dashed grey lines denote asynchronous trials. Shaded bands represent 95 % confidence intervals. **c.** Mean pupil dilation averaged across 12 trials, shown separately for the systole session (left) and diastole session (right). Boxplots depict synchronous (pink for systole, blue for diastole) and asynchronous (grey) conditions; individual data points are overlaid. **d.** Model-predicted looking time as a function of pupil dilation. Pink line represents the systole session; blue line the diastole session. Shaded bands represent 95 % confidence intervals.

Thereafter, we averaged each infant’s pupil change over this 4 s window and entered these values into separate LMMs for the systole and diastole sessions, using the same model structure as that for looking time. In the systole session, the full model outperformed a null model (*χ²*(3) = 16.77, *p* < 0.001) and a main-effects model (*χ²*(1) = 4.80, *p* = 0.028). Crucially, there was a significant synchrony × age interaction (β = 0.040, *t*(364.8) = 2.19, *p* = 0.029; Table 2). A follow-up contrast confirmed that pupil dilation to synchronous stimuli increased with age, whereas dilation to asynchronous stimuli remained relatively stable (*t*(364) = –2.19, *p* = 0.029; Fig. 3B, left panel). Separate LMMs within each age-group showed the synchrony effect only in younger infants (β = –0.071, *t*(184.9) = –2.68, *p* = 0.0080) and not in older infants (β = 0.015, *t*(178.2) = 0.61, *p* = 0.54). The raw distributions of individual mean pupil changes across trials are shown in Fig. 3C, left panel.

**Table 2.**
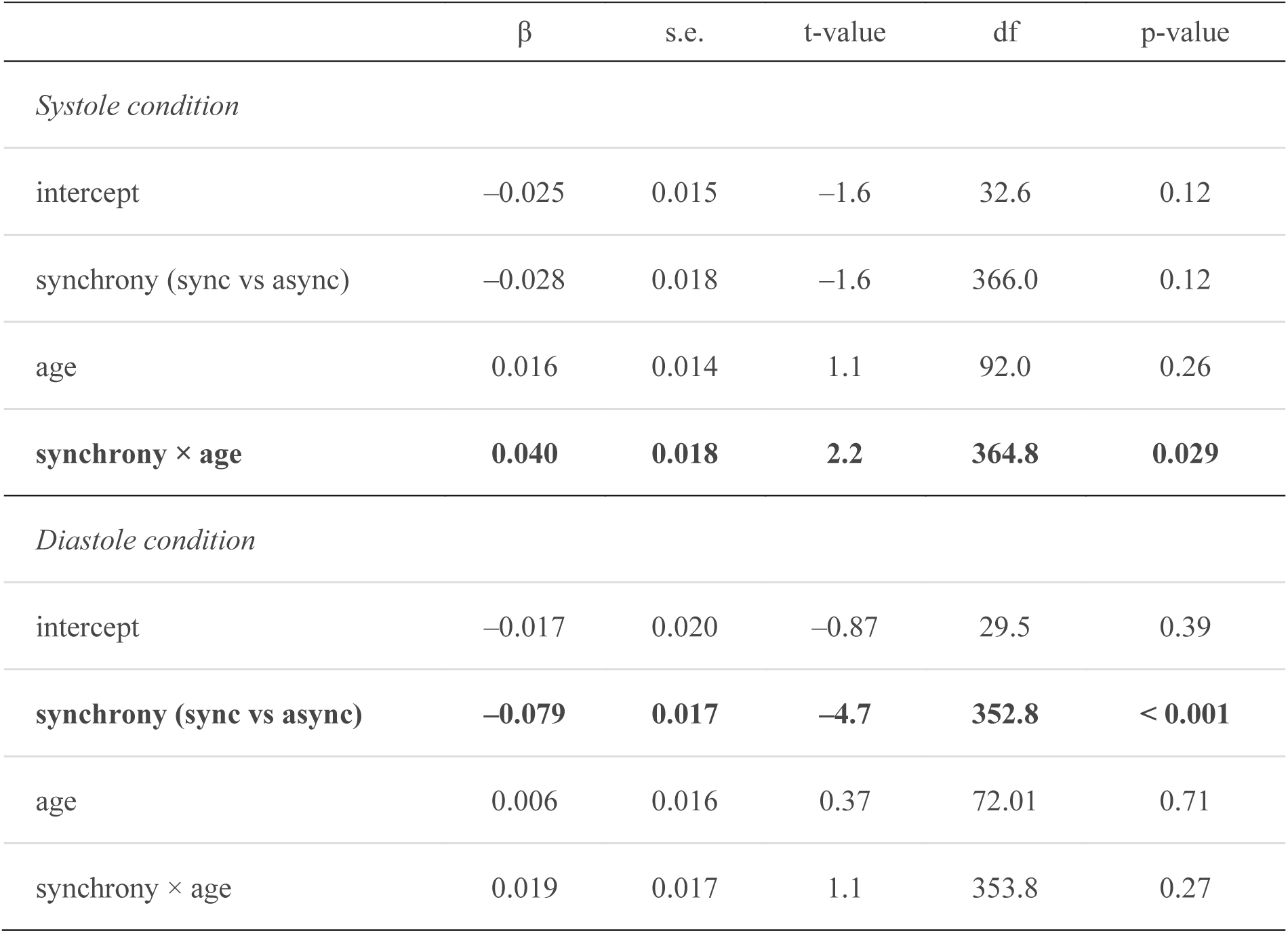
Results of the linear mixed model predicting pupil change.

| | $\beta$ | s.e. | t-value | df | p-value |
| --- | --- | --- | --- | --- | --- |
| *Systole condition* |  |  |  |  |  |
| intercept | -0.025 | 0.015 | -1.6 | 32.6 | 0.12 |
| synchrony (sync vs async) | -0.028 | 0.018 | -1.6 | 366.0 | 0.12 |
| age | 0.016 | 0.014 | 1.1 | 92.0 | 0.26 |
| **synchrony × age** | **0.040** | **0.018** | **2.2** | **364.8** | **0.029** |
| *Diastole condition* |  |  |  |  |  |
| intercept | -0.017 | 0.020 | -0.87 | 29.5 | 0.39 |
| **synchrony (sync vs async)** | **-0.079** | **0.017** | **-4.7** | **352.8** | **< 0.001** |
| age | 0.006 | 0.016 | 0.37 | 72.01 | 0.71 |
| synchrony × age | 0.019 | 0.017 | 1.1 | 353.8 | 0.27 |

In the diastole session, although the full model outperformed the null model (*χ²*(3) = 23.8, *p* < 0.001), it did not improve in the main-effects model (*χ²*(1) = 1.23, *p* = 0.27). Accordingly, only a main effect of synchrony was significant (*β* = –0.079, *t*(353.8) = –4.70, *p* < 0.001, Table 2), indicating reduced dilation for diastole-locked synchronous stimuli across ages (Fig. 3B–C, right). Together, these results suggest that synchronous stimuli initially suppress pupil dilation—presumably via baroreflex mechanisms (see Discussion)—but this inhibition weakens with age when stimuli fall within the baroreceptor-active systolic window, leading older infants to show enhanced dilation to synchronous stimuli. Data on fast versus slow asynchronous rhythms are presented in Table S2.

Finally, we tested whether pupil dilation predicts looking time. In the systole session, an LMM (looking time ∼ pupil + age + pupil:age + (1|participant) + (1|trial)) revealed that pupil dilation was the only significant predictor (*β* = –2.00, *t*(363.6) = –2.21, *p* = 0.028). By contrast, no effects emerged in the diastole session (Table 3; Fig. 3D). Thus, in the systole session, larger pupil dilation accounts for why older infants look away sooner from synchronous than asynchronous stimuli, linking the physiological and behavioural indices of cardiac-exteroceptive integration.

**Table 3.** Results of the linear mixed model predicting looking time from pupil change.

| | $\beta$ | s.e. | t-value | df | p-value |
| --- | --- | --- | --- | --- | --- |
| *Systole condition* |  |  |  |  |  |
| intercept | 6.4 | 0.83 | 7.7 | 16.2 | < 0.001 |
| **pupil-change** | **-2.00** | **0.91** | **-2.2** | **363.6** | **0.028** |
| age | -0.51 | 0.39 | -1.3 | 34.8 | 0.21 |
| pupil-change x age | 0.32 | 1.0 | 0.32 | 367.3 | 0.675 |
| *Diastole condition* |  |  |  |  |  |
| intercept | 7.5 | 0.83 | 9.0 | 20.4 | < 0.001 |
| pupil-change | 0.33 | 1.2 | 0.28 | 372.7 | 0.78 |
| age | -0.63 | 0.49 | -1.3 | 35.4 | 0.21 |
| pupil-change x age | 0.76 | 1.3 | 0.58 | 368.9 | 0.56 |

### Association with autonomic function

To relate infants’ looking behaviour in the systole session to autonomic function, we calculated a looking-time bias index (LBI) for each pair of trials—for example, trials 1–2, 3–4, 5–6, …—in which one trial was synchronous and the other asynchronous, as follows: 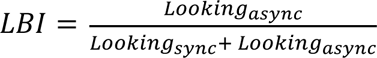. A larger LBI, thus, indicates a stronger preference for asynchronous stimuli. We then fitted an LMM predicting LBI from age, resting-state heart rate (HR), high-frequency (HF) HRV (vagal index), and low-frequency (LF) HRV (overall autonomic index), with a random intercept for trial-pair (LBI ∼ age + HR + HF + LF + (1|trial-pair)). All continuous predictors were z-standardised. This model improved fit over the one with only age (*χ²*(3) = 8.10, *p* = 0.044). Both age (*β* = 0.025, *t*(664) = 2.62, *p* = 0.0090), and LF (*β* = 0.030, *t*(664) = 2.66, *p* = 0.0081) emerged as significant predictors (Table 4). These results suggest that the developmental shift in the LBI is partly driven by autonomic maturation; thus, bolstering the index’s validity as a marker of interoceptive processing in infancy.

**Table 4.** Results of the linear mixed model predicting Looking-Time Bias (LBI) from HRV.

| | $\beta$ | s.e. | t-value | df | p-value |
| --- | --- | --- | --- | --- | --- |
| intercept | 0.52 | 0.0088 | 58.9 | 664 | < 0.001 |
| **age** | **0.025** | **0.010** | **2.6** | **664** | **0.009** |
| heart rate | 0.0084 | 0.012 | 0.73 | 664 | 0.47 |
| HRV-HF | -0.0053 | 0.012 | -0.43 | 664 | 0.766 |
| **HRV-LF** | **0.030** | **0.011** | **2.66** | **664** | **0.008** |

## Discussion

The continuous integration of exteroceptive and interoceptive signals is fundamental for evaluating sensory inputs and guiding adaptive behaviours (Engelen et al., 2023). This sophisticated interplay underpins the formation of a coherent sense of self, and facilitates effective responses to changing circumstances. However, the developmental origins of such integration remain poorly understood. In the present study, we examined its early developmental trajectory, showing how infants begin to combine internal bodily states with external sensory cues to scaffold their perception, cognition, and action.

Using a refined iBEATs paradigm, and building upon the pioneering study by Maister et al. (2017), we found that infants develop the ability to discriminate cardiac-exteroceptive synchrony between 3 and 8 months of age, as indexed by a preference for asynchronous over synchronous stimuli, matched to their own heartbeats. Notably, this effect was observed only when the synchronous stimulus was aligned with the cardiac systole—the baroreceptor-active phase during which the central representations of cardiac signals are most prominent. In contrast, no such effect was observed when the stimuli were synchronised with the timing of the cardiac R-wave—a phase marked by electrical depolarisation but minimal baroreceptor input (i.e. diastole). This pattern strongly suggests that observed behavioural responses are driven by the central integration of cardiac signals with exteroceptive information, rather than by simple temporal matching of heartbeat rhythms. Furthermore, our results effectively ruled out the possibility that Donaghy et al. (2025) pointed outinfants’ preference for asynchronous stimuli could be driven by a novelty bias due to greater variability in the asynchronous stimuli (± 10% of the previous trial). If this were the case, infants should have preferred asynchronous stimuli equally in both the systole and diastole sessions. The absence of such a pattern supports our interpretation that the observed preference reflects genuine interoceptive–exteroceptive integration.

We observed a developmental shift in behavioural responses between 3- and 8-month-old infants. Specifically, only the older group of infants aged 6–8 months demonstrated a clear preference for asynchronous over synchronous stimuli in the systole session. This aligns with the neurodevelopmental evidence indicating that the anterior insula and anterior cingulate cortices—key hubs of the salience network, that functionally coordinate with the default mode network and are thought to underpin the integration of interoceptive and exteroceptive information by enabling long-range functional connectivity across cortical regions (Feldman Barrett & Simmons, 2015)—are absent in new-borns, but undergo rapid maturation by 6 months of age (Gao, Alcauter, Elton, et al., 2015; Gao, Alcauter, Smith, et al., 2015; van den Heuvel et al., 2012, 2013). The maturation of these networks likely enables infants to integrate cardiac-exteroceptive signals, guiding their behaviour, as reflected in the looking preferences observed in this study. Importantly, these networks are also proposed to support predictive brain functions (Feldman Barrett & Kyle Simmons, 2015). As discussed below, the behavioural patterns observed in our study can be meaningfully interpreted within this computational framework.

Several previous studies employing iBEATs have interpreted infants’ stimulus preferences within a familiarity-novelty dichotomy (Imafuku et al., 2023; Tünte et al., 2025). In these studies, synchronous stimuli are assumed to be more familiar and easier to process, although the reported findings diverge: Imafuku et al. (2023) reported a novelty preference, whereas Tünte et al. (2025) found evidence for a familiarity preference. While these interpretations offer valuable insights, we argue that exteroceptive stimuli that co-vary with internal signals are not inherently ‘familiar’ in the traditional sense based solely on repeated exposure. Moreover, the absence of a preference effect in the diastole—despite the stimuli being identical in rhythmic structure to those used in the systole session challenges a purely familiarity-based account. Instead, we propose that the observed behavioural effects are better explained by dynamic computational processes grounded in predictive-coding mechanisms. Recent theoretical work, supported by empirical research, suggests that infants’ looking behaviour reflects sophisticated statistical inferences, driven by a Bayesian evaluation of how well observed data support competing hypotheses (Sim & Xu, 2019). These evaluations guide exploratory behaviours such as prolonged looking, to refine internal perceptual models. Within the predictive-coding framework, such hypotheses are conceptualised as ‘predictions’ generated by the brain, and continuously tested through sensory sampling. When incoming information deviates from these predictions, a ‘prediction-error’ is produced, prompting model updating (Clark, 2013; Friston, 2009; Hohwy, 2013; Lee & Mumford, 2003). It has further been proposed that coordinated predictions constitute the basis of a ‘concept’—a structured internal model of expected sensory regularities—and that concept formation involves encoding and refining such predictive representations (Atzil et al., 2018). Crucially, temporal contingencies between interoceptive and exteroceptive signals have been emphasised as a key factor for constructing such concepts, and forming accurate predictions (Atzil et al., 2018; Engelen et al., 2023; Feldman Barrett & Simmons, 2015; Seth, 2013). Atzil et al. (2018) specifically identify 6 months as a key developmental milestone for the emergence of conceptual abilities, underpinned by the maturation of the aforementioned neural networks. In the context of this study, we propose that for older infants whose neural networks have matured, exteroceptive signals, when effectively integrated with cardiac signals, facilitate the rapid construction of perceptual models of the stimuli, enabling accurate predictions about incoming sensory input. When audiovisual stimuli are continuously presented in synchrony with internal cardiac signals, they closely align with these predictions. Consequently, there is minimal need for model updating or further sensory sampling, leading to shorter looking times. In contrast, asynchronous stimuli disrupt this alignment, triggering larger prediction errors and necessitating model revision, which results in longer looking times. Notably, this effect was specific to the systole session. In the diastole session, the reduced salience of the cardiac signals may have impaired the efficiency of such integration. Our pupillary data provide further converging support for this interpretation.

In the systole session, we observed a developmental increase in pupil dilation in response to synchronous stimuli, whereas no age-related change was found in the diastole session. Although no prior studies have directly examined pupillary responses during interoceptive–exteroceptive integration, research on audiovisual multisensory integration has shown greater pupil dilation for multimodal compared to unimodal presentations, with responses to multimodal stimuli often exceeding the sum of unimodal responses (Rigato et al., 2016; Wang et al., 2017). While the pupil diameter has been linked to a broad range of cognitive and affective processes, including attention, arousal, surprise, and learning (Viglione et al., 2023), recent theoretical accounts converge on the idea that pupil dilation reflects information gain—the degree to which sensory input updates internal models. This has been formalised as the Kullback–Leibler divergence between prior and posterior beliefs (Zénon, 2019), or as a related (but not equivalent) concept within predictive coding frameworks, namely, precision-weighted prediction error (Harris et al., 2022). Neurophysiologically, pupil dilation is mediated by phasic noradrenaline release from the locus coeruleus, which is thought to amplify the influence of sensory input on perception, and accelerate belief updating. From this perspective, the developmental increase in pupil dilation in response to systole-locked synchronous stimuli observed in this study may reflect the process by which these stimuli generate prediction-error signals to higher-order cortical areas and trigger perceptual model updating—specifically in infants whose predictive systems supporting interoceptive-exteroceptive integration have matured. Asynchronous stimuli, by violating temporal predictions, also engage in model-updating processes, and thus elicit pupil dilation. This may explain why older infants’ pupil responses to synchronous and asynchronous stimuli were comparable—at least within the first 4 s of the first 12 trials, during which belief updating for the synchronous stimuli is likely to be most pronounced. By contrast, synchronous stimuli in the diastole session that do not induce central integration may have produced minimal pupil dilation.

However, pupil responses to diastole-locked synchronous stimuli may require more cautious interpretation. Notably, in younger infants whose predictive systems are presumably still immature—pupil size was consistently smaller in response to synchronous stimuli—systole- or diastole-locked—than to asynchronous stimuli. Although the underlying mechanisms remain unclear, we propose that this pattern may reflect low-level physiological processes, specifically baroreflex-mediated responses. Pupil size is known to fluctuate spontaneously, even in the absence of external stimulation. Recent studies have shown that these spontaneous fluctuations are modulated by baroreflex-related autonomic activity (Bär et al., 2009; Calcagnini et al., 2005), primarily attributed to parasympathetic influence. Baroreflex activation induces pupil constriction via the parasympathetic pathways innervating the sphincter pupillae muscle—distinct from the sympathetic control of the dilator pupillae muscle, which governs dilation responses associated with the cognitive and arousal processes discussed above. It is, therefore, possible that audiovisual stimuli presented in synchrony with the cardiac cycle may coincide with the phases of increased parasympathetic tone, thereby inhibiting dilation responses. This inhibitory effect may dominate in younger infants, as their higher-level cognitive systems for interoceptive–exteroceptive integration are likely still underdeveloped. In older infants, however, we observed a dissociation: pupil constriction was evident in the diastole condition—likely reflecting parasympathetic dominance—whereas in the systole condition, cognitively driven dilation responses appear to override this reflexive constriction. Although these interpretations remain speculative, the results from an ad hoc adult experiment (n = 11) closely resembled those observed in older infants (Fig. S1), suggesting that the inhibition of pupil dilation to R-wave-locked stimuli is a robust phenomenon in adulthood, and that the responses in the older group reflect the emergence of mature, adult-like mechanisms.

We observed that infants’ growing preference for asynchronous over synchronous stimuli was predicted by the LF power of their 5-min resting HRV, whereas HF power showed no such association. Originally, we had hypothesized—based on adult work linking vagal tone and interoceptive processing (Blickle et al., 2024; Lischke et al., 2020; Paciorek & Skora, 2020; Richter et al., 2021; Rominger et al., 2021; Vabba et al., 2023; Villani et al., 2019)—that HF power would predict this behavioural shift. However, in infancy, rapid and irregular breathing may undermine HF power’s validity as a parasympathetic index (Chiera et al., 2020; Latremouille et al., 2021). Infant studies have applied a variety of HF bands owing to the lack of a standardised range (Latremouille et al., 2021). We followed the 0.24–1.04 Hz guideline proposed by Quintana et al. (2016), but how accurately this range captures infant vagal function remains unclear. By contrast, LF power reflects a mixture of baroreflex-mediated vagal modulation and sympathetic influences, and is generally considered an index of overall autonomic nervous system (ANS) maturation in infants. Although it is difficult to disentangle the specific contribution of sympathetic and parasympathetic branches to LF power in infancy, we assume that it more strongly reflects vagal maturation, considering that the mammalian vagus is only partially myelinated at birth and continues to develop postnatally—with the steepest increase occurring between approximately 30–32 weeks of gestation, and 6 months after birth (Porges & Furman, 2011). In contrast, the sympathetic nervous system is thought to develop during foetal life, and is relatively more mature at birth (Porges & Furman, 2011). Vagal maturation enables more effective visceral regulation, which in turn, facilitates better behavioural regulation in infancy (Porges & Furman, 2011). Taken together, our findings suggest that autonomic maturation—and possibly vagal development in particular—underpins infants’ emerging capacity to integrate internal bodily signals with external sensory events, and regulate behaviours based on the representation. Although further validation is warranted, the observed association between ANS function and looking-time behaviour provides converging evidence that the iBEATs task captures infants’ interoceptive processing.

Additionally, we observed that maternal interoceptive sensibility, as assessed by the Multidimensional Assessment of Interoceptive Awareness (MAIA), predicted infants’ behavioural sensitivity to cardiac–exteroceptive synchrony (see Table S3 and Supplementary Results). While this finding is preliminary and insufficient to draw firm conclusions about how maternal traits influence the development of interoceptive capacities in infants, it offers tentative support for theoretical frameworks proposing that early interoceptive-exteroceptive integration emerges primarily within the caregiving context, in which caregiver interoceptive inference plays a key role (Fotopoulou & Tsakiris, 2017; see Supplementary Discussion). Further investigation is warranted to explore this possibility more systematically.

Overall, this study provides the first evidence that the ability to integrate cardiac and exteroceptive signals emerges during the first year of life, and that this process is influenced by autonomic maturation. The behavioural and physiological findings are well explained within the predictive coding framework. By offering a multidimensional account of how cardiac–exteroceptive integration develops in infancy, this study lays the groundwork for future investigations into the ontogeny of interoceptive processing. Moreover, its findings support the validity of iBEATs as a sensitive tool for probing early interoceptive–exteroceptive integration, but only when stimuli are time-locked to cardiac systole rather than to the R-wave. Future studies should further examine whether vagal pathways are causally involved in this process, and how central neural systems contribute to the integration of interoceptive and exteroceptive signals. It also remains to be determined whether sensitivity measured by the iBEATs task predicts individual differences in broader behavioural outcomes, such as emotion regulation, attention, and cognitive functioning. This modified iBEATs paradigm, therefore, offers a promising avenue for advancing our understanding of interoceptive development in early life.

## Methods

### Participants

The final sample comprised 36 infants (23 girls) and their mothers, whose ages ranged from 3–8 months (mean age = 179.1 days, SD 45.5, range: 100–262 days). An additional 12 infants were tested, but excluded from the analyses owing to fussiness (n = 2), abnormal ECG waveforms (n = 1), failure in stimulus presentation (n = 1), insufficient usable eye-tracking data (n = 7), or strong side bias in looking behaviour (n = 1). A simulation-based power analysis was conducted using the *simr* package in R (Green & MacLeod, 2016). Assuming a moderate effect size (β = 0.50) for the interaction term of *synchrony* × *age* in the model (lookingTime ∼ synchrony + age + synchrony:age + (1|id) + (1|trial), the analysis yielded an estimated power of 80.0% (95% CI: 70.8%–87.3%) with a sample size of 36 participants and α = .05, based on 100 simulations using Satterthwaite approximation for degrees of freedom. The participants were recruited from the database of Unicharm Co., Japan. This study was approved by the Ethics Committee of Nagoya University’s Department of Psychological and Cognitive Sciences (approval code: NUPSY-220421-G-01). Written informed consent was obtained from all the participating parents on behalf of themselves and their infants.

### Apparatus

Infants’ heartbeats were recorded using ECG100C biometric amplifiers connected to a BIOPAC MP160 system (BIOPAC Systems, Inc.). Two distinct ECG amplifiers were used. The first amplifier captured raw ECG waveforms and was configured with a 1.0 Hz high-pass filter and a 35 Hz low-pass filter applied online. The second amplifier was dedicated to R-peak detection and operated in the build-in ‘R-wave detector mode’, which outputs a smoothed pulse at each R-wave occurrence, even under conditions with signal artefacts. Although this mode introduces a delay of approximately 50 ms owing to filtering, it was chosen to ensure robust online R-wave detection for trigger generation, as infants’ ECG signals often fluctuate owing to body movements. The sampling rate was set at 1000 Hz, and a 60 Hz notch filter was applied online. The R-waves were detected online using a digital trigger unit (DTU100C), that received input from the second ECG amplifier via the MP160 system. A digital TTL trigger was generated when the analogue input exceeded a predefined threshold, and was sent to a PC via an external digital I/O module (National Instruments, Ltd.).

The infants’ eye gaze was recorded using a Tobii Pro Spectrum eye tracker (Tobii AB), operating at 60 Hz. Eye-tracking data were transmitted in real time to a PC using the Tobii Pro SDK. Stimulus presentation was controlled on the same PC, which integrated inputs from both the ECG and eye-tracking systems. All task controls and data integration were managed using a custom script written in Psychtoolbox-3 (Brainard, 1997; Pelli, 1997; Kleiner et al., 2007), within MATLAB (MathWorks, Inc.).

### Stimuli

We presented infants with an audiovisual animated stimulus that pulsed either in sync or out of sync with their heartbeat. The visual component consisted of Greebles (Gauthier & Tarr, 2002), which are novel, unfamiliar shapes, that nonetheless share certain structural constraints with human faces. We selected 30 original Greeble shapes and altered their surface colours to ensure that each trial used a distinct stimulus, thereby minimising the influence of shape or colour preference on the infants’ looking behaviour. If an infant completed more than 30 trials, the same set of 30 stimuli were reused in a random order for an additional block of 30 trials. To enhance the infants’ attention, each shape was augmented with cartoon-like eyes, following prior studies (Imafuku et al., 2023; Maister et al., 2017; Weijs et al., 2023; Figure 1A). The auditory component was identical across all trials and consisted of a 100 ms tone obtained from a free online sound library (OtoLogic). Visually, the stimuli rhythmically expanded and contracted to create the impression of movement. Before the onset of each animated stimulus, we presented a pixel-shuffled mask image—created from the corresponding stimulus using Adobe Illustrator (Adobe Inc.). The mask matched the animated stimulus in terms of mean luminance, thereby minimising pupil response artefacts related to luminance changes. At the beginning of the task and during the inter-trial intervals, we showed infant-friendly video clips from Baby Mozart (Walt Disney Video) to maintain attention.

### Procedure

Each family visited the laboratory on two separate days, spaced a week apart: one for the *systole* session and the other for the *diastole* session. After the parents provided informed consent, five disposable electrodes were attached to the infants’ chests and abdomens—two for recording raw ECG signals (connected to the first amplifier), two for detecting R-waves (connected to the second amplifier), and one ground electrode. Infants were then seated on their parents’ laps, and an eye-tracking calibration procedure was conducted, to confirm that their gaze could be reliably tracked at the left, centre, and right positions on the screen where the stimuli would be presented. Once both ECG signal acquisition and eye-tracking accuracy were confirmed, the iBEATs task was introduced.

At the beginning of the task, a 20 s infant-friendly animation from Baby Morzart (Walt Disney Video) was shown. During this period, the infant’s average RR interval was calculated and used to control stimulus presentation in subsequent trials (e.g. detected triggers that deviated substantially from this baseline were ignored in real time). Each trial began with a short fixation clip (minimum 1 s), followed by a pixel-shuffled mask stimulus (at least 0.5 s). Gaze was monitored during both phases, and once the infants fixated on the area, the trial proceeded to the main stimulus presentation. In each trial, the infants viewed an audiovisual stimulus that rhythmically expanded and contracted, either synchronously with their own heartbeat (systole or diastole) or asynchronously. In the synchronous trial, the expanded image was time-locked to either + 250–300 or + 0–50 ms R-waves for the systole and diastole sessions, respectively (see Fig. 1A for an example of the stimulus-timing distribution). The expanded image was displayed for 100 ms, followed by the contracted image, which remained on screen until the next trigger. A 100 ms auditory tone was presented simultaneously with the expanded image. In the asynchronous trials, the heartbeat rhythm was taken from another infant in a previous study (Suga et al., 2019). Specifically, a sample whose RR interval closely matched that of the current infant’s most recent trial was selected. A different segment of this sample was used for each asynchronous trial. The RR intervals were further adjusted to be 10 % faster or slower than the infant’s mean heart rate, calculated from the previous trial (or from the initial 20 s animation for the first trial). Synchronous and asynchronous trials were alternated, and the stimulus was presented on the screen’s left or right side. Each trial lasted up to 20 s, as long as the infant maintained fixation. If the infant consecutively looked away for more than 3 s, the trial ended, and the next one began. The task comprised up to 60 trials, but the session was terminated earlier if the infant became tired or fussy. The stimulus type (synchronous or asynchronous), its position (left or right), and session order (systole or diastole) were counterbalanced across participants. However, these parameters were held constant within each infant across the two sessions. For example, if an infant began the diastole session on the first day with a synchronous stimulus on the left side, the same pattern was followed in the systole session on the second day.

After the iBEATs task, the infant’s resting ECG was recorded for 5 minutes while the infant lay quietly.

### Questionnaires

Parents were asked to complete a set of questionnaires either before or after their laboratory visits, using online forms, which was part of a larger investigation involving infants and their parents. Among these questionnaires, the Japanese version of MAIA(MAIA-J); (Mehling et al., 2012; Shoji et al., 2018) was used for supplementary analyses in this study.

### Data analyses

#### ECG processing and check for real-time synchronisation

First, we verified whether stimulus presentation was successfully time-locked to the targeted cardiac timing. Using a custom-written script in MATLAB (MathWorks, Inc.), we segmented the raw ECG signals recorded by the first amplifier into individual trials, and detected the time points of R-peaks. For each synchronous trial, we calculated the timing of stimulus presentation relative to the nearest R-peak. Trials in which the stimulus was not presented within the targeted window (R + 250–300 ms in the systole session; R + 0–50 ms in the diastole session) were identified based on the SD thresholds and visual inspection. Synchronisation failures included: (1) stimulus presentation outside the specified time window; (2) stimulus omission due to missed R-wave detection; or (3) multiple stimuli presented within a single cardiac cycle owing to signal artefacts. Any trial exhibiting such errors was excluded from further analysis. In asynchronous trials, because the stimuli were presented based on preloaded heartbeat data, real-time synchronisation failures did not occur.

#### Looking time

Eye-tracking data, including fixation coordinates and pupil size at each time point (sampled at 60 Hz), were recorded using a Tobii Pro Spectrum eye tracker and stored in MATLAB. During the task, based on the average coordinates of the left and right eye, the system continuously analysed in real time whether the infants’ point of gaze fell within the area of the presented stimulus at each sampling point to dynamically control the task. For the offline analysis, these data were used to calculate the total duration that the infant looked at the stimulus during each trial. The resulting trial-level looking times were then entered into a LMM, with trial-level labels indicating session type (systole versus diastole) and stimulus type (synchronous versus asynchronous).

#### Pupil

Raw pupil data output from the Tobii Pro SDK were first segmented into individual trials, beginning 200 ms prior to stimulus onset (during the mask presentation) and continuing until the end of each trial. Artefacts were detected by identifying data points that exceeded 3 SDs from the mean pupil size or mean velocity; these points were replaced with NaNs. As reliable pupil measurements from both eyes, were not always available, data from the eye with the higher data retention rate was used. For each trial, time-series changes in pupil size were calculated by subtracting the mean pupil size during the 200 ms pre-stimulus baseline (mask presentation) from the pupil size during the stimulus animation. As pupil data could only be recorded when the infant was fixating on the screen, and fixation declined over time within and across trials, our analysis was restricted to the first 4 s following stimulus onset in the first 12 trials (6 synchronous, 6 asynchronous). This time window was selected because more than 70 % of pupil samples were successfully tracked during this period. For visualisation purposes, a 10-point moving average was applied to smoothen the time-series pupil data. For statistical analysis, the average pupil size over the 4 s window was entered into the LMM.

#### Heart Rate Variability

The infants’ 5-minute resting ECG data, recorded after the task on the first day were used to calculate HRV. For infants whose resting data could not be collected due to fussiness, data from the second day were used instead (n = 5). R-peaks were first detected using a custom-written MATLAB script, and a time series of interbeat intervals was constructed. Subsequently, these data were imported into the RHRV package in R (Garcia Martinez et al., 2017). The minimum and maximum heart rates were set to 60 and 180 bpm, respectively, and detected outliers were adjusted using interpolation. Frequency-domain analysis was performed using Fourier methods. The LF and HF bands were defined as 0.04–0.24 and 0.24–1.0 Hz, respectively, in accordance with the guidelines provided by Quintana et al. (2016).

#### Linear Mixed Models

Model fitting was conducted using the lmer function from the lme4 package in R (Bates et al., 2015). Estimated marginal means and pairwise comparisons were calculated using the emmeans package (Lenth et al., 2021).

## Acknowledgement

We extend our sincere gratitude to Yosuke Naruto and Sakina Ubukata for their invaluable assistance in data collection. We also thank Dr. Lara Maister for generously sharing insights from her previous work. We are profoundly grateful to all the infants and their families for their cooperation and contribution to this study. Support for this study was provided by JST CREST to HO (#JPMJCR21P1) and JSPS Kakenhi to TI (#22H03494; #23K24751).

## Author contributions

AS, TI and HO led the project. MK and TI collected the preliminary data, and AS collected the main data. TI analysed the data and drafted the manuscript. All authors contributed to the study’s design, interpretation of results, and manuscript revision, as well as to its final version’s review and approval.

## References

Atzil, S., Gao, W., Fradkin, I., & Barrett, L. F. (2018). Growing a social brain. In Nature Human Behaviour (Vol. 2, Issue 9, pp. 624–636). Nature Publishing Group. 10.1038/s41562-018-0384-6

Bär, K.-J., Schulz, S., Koschke, M., Harzendorf, C., Gayde, S., Berg, W., Voss, A., Yeragani, V. K., & Boettger, M. K. (2009). Correlations between the autonomic modulation of heart rate, blood pressure and the pupillary light reflex in healthy subjects. Journal of the Neurological Sciences, 279(1–2), 9–13.

Bates, D., Mächler, M., Bolker, B., & Walker, S. (2015). Fitting linear mixed-effects models Usinglme4. Journal of Statistical Software, 67(1), 1–48.

Blickle, M., Klüpfel, C., Homola, G. A., Gamer, M., Herrmann, M. J., Störk, S., Gelbrich, G., Heuschmann, P. U., Deckert, J., Pham, M., & Menke, A. (2024). Heart rate variability, interoceptive accuracy and functional connectivity in middle-aged and older patients with depression. Journal of Psychiatric Research, 170, 122–129.

Brener, J., & Kluvitse, C. (1988). Heartbeat detection: judgments of the simultaneity of external stimuli and heartbeats. Psychophysiology, 25(5), 554–561.

Brown, G. L., & Eccles, J. C. (1934). The action of a single vagal volley on the rhythm of the heart beat. The Journal of Physiology, 82(2), 211–241.

Calcagnini, G., Giovannelli, P., Censi, F., Bartolini, P., & Barbaro, V. (2005). Baroreceptor-sensitive fluctuations of heart rate and pupil diameter. 2001 Conference Proceedings of the 23rd Annual International Conference of the IEEE Engineering in Medicine and Biology Society, 1, 600–603 vol.1.

Charbonneau, J. A., Maister, L., Tsakiris, M., & Bliss-Moreau, E. (2022). Rhesus monkeys have an interoceptive sense of their beating hearts. Proceedings of the National Academy of Sciences of the United States of America, 119(16), e2119868119.

Chiera, M., Cerritelli, F., Casini, A., Barsotti, N., Boschiero, D., Cavigioli, F., Corti, C. G., & Manzotti, A. (2020). Heart rate variability in the perinatal period: A critical and conceptual review. Frontiers in Neuroscience, 14, 561186.

Clark, A. (2013). Whatever next? Predictive brains, situated agents, and the future of cognitive science. The Behavioral and Brain Sciences, 36(3), 181–204.

Donaghy, R., Lisi, M., Shinskey, J., & Murphy, J. (2025). Measuring cardiac interoceptive accuracy in infancy: Lessons from the adult literature. Psychophysiology, 62(3), e70041.

Engelen, T., Solcà, M., & Tallon-Baudry, C. (2023). Interoceptive rhythms in the brain. Nature Neuroscience, 26(10), 1670–1684.

Feldman Barrett, L., & Kyle Simmons, W. (2015). Interoceptive predictions in the brain. Nature Reviews. Neuroscience, 16, 419–429.

Fotopoulou, A., & Tsakiris, M. (2017). Mentalizing homeostasis: the social origins of interoceptive inference. Neuropsychoanalysis, 4145(June), 1–9.

Friston, K. (2009). The free-energy principle: a rough guide to the brain? Trends in Cognitive Sciences, 13(7), 293–301.

Gao, W., Alcauter, S., Elton, A., Hernandez-Castillo, C. R., Smith, J. K., Ramirez, J., & Lin, W. (2015). Functional network development during the first year: Relative sequence and socioeconomic correlations. Cerebral Cortex (New York, N.Y.: 1991), 25(9), 2919–2928.

Gao, W., Alcauter, S., Smith, J. K., Gilmore, J. H., & Lin, W. (2015). Development of human brain cortical network architecture during infancy. Brain Structure & Function, 220(2), 1173–1186.

Gao, W., Zhu, H., Giovanello, K. S., Smith, J. K., Shen, D., Gilmore, J. H., & Lin, W. (2009). Evidence on the emergence of the brain’s default network from 2-week-old to 2-year-old healthy pediatric subjects. Proceedings of the National Academy of Sciences of the United States of America, 106(16), 6790–6795.

Garcia Martinez, C. A., Otero Quintana, A., Vila, X. A., Tourino, M. J. L., Rodriguez-Linares, L., Rodriguez Presedo, J. M., & Mendez Penin, A. J. (2017). Heart rate variability analysis with the R package RHRV [PDF]. Springer International Publishing.

Garfinkel, S. N., & Critchley, H. D. (2016). Threat and the body: How the heart supports fear processing. Trends in Cognitive Sciences, 20(1), 34–46.

Gauthier, I., & Tarr, M. J. (2002). Unraveling mechanisms for expert object recognition: Bridging brain activity and behavior. Journal of Experimental Psychology. Human Perception and Performance, 28(2), 431–446.

Green, P., & MacLeod, C. J. (2016). SIMR: an R package for power analysis of generalized linear mixed models by simulation. Methods in Ecology and Evolution, 7(4), 493–498.

Harris, D. J., Arthur, T., Vine, S. J., Liu, J., Abd Rahman, H. R., Han, F., & Wilson, M. R. (2022). Task-evoked pupillary responses track precision-weighted prediction errors and learning rate during interceptive visuomotor actions. Scientific Reports, 12(1), 22098.

Hickman, L., Seyedsalehi, A., Cook, J. L., Bird, G., & Murphy, J. (2020). The relationship between heartbeat counting and heartbeat discrimination: A meta-analysis. Biological Psychology, 156, 107949.

Hohwy, J. (2013). The Predictive Mind. Oxford University Press.

Imafuku, M., Yoshimoto, H., & Hiraki, K. (2023). Infants’ interoception is associated with eye contact in dyadic social interactions. Scientific Reports, 13(1), 9520.

Izard, C. E., Porges, S. W., Simons, R. F., Haynes, O. M., Hyde, C., Parisi, M., & Cohen, B. (1991). Infant cardiac activity: Developmental changes and relations with attachment. Developmental Psychology, 27(3), 432–439.

Latremouille, S., Lam, J., Shalish, W., & Sant’Anna, G. (2021). Neonatal heart rate variability: a contemporary scoping review of analysis methods and clinical applications. BMJ Open, 11(12), e055209.

Lee, T. S., & Mumford, D. (2003). Hierarchical Bayesian inference in the visual cortex. Journal of the Optical Society of America. A, Optics, Image Science, and Vision, 20(7), 1434–1448.

Lenth, R. V., Banfai, B., Bolker, B., Buerkner, P., Giné-Vázquez, I., Herve, M., Jung, M., Love, J., Miguez, F., Piaskowski, J., Riebl, H., & Singmann, H. (2021). emmeans: Estimated marginal means, aka least-squares means. https://cran.r-project.org/package=emmeans.

Levy, M. N., & Martin, P. J. (1984). Neural control of the heart. In Physiology and Pathophysiology of the Heart (pp. 337–354). Springer US.

Lischke, A., Pahnke, R., Mau-Moeller, A., & Weippert, M. (2020). Heart rate variability modulates interoceptive accuracy. Frontiers in Neuroscience, 14, 612445.

Maister, L., Tang, T., & Tsakiris, M. (2017). Neurobehavioral evidence of interoceptive sensitivity in early infancy. ELife, 6, 1–12.

Mehling, W. E., Price, C., Daubenmier, J. J., Acree, M., Bartmess, E., & Stewart, A. (2012). The Multidimensional Assessment of Interoceptive Awareness (MAIA). PloS One, 7(11), e48230.

Paciorek, A., & Skora, L. (2020). Vagus Nerve Stimulation as a gateway to interoception. Frontiers in Psychology, 11, 1659.

Pollatos, O., & Schandry, R. (2004). Accuracy of heartbeat perception is reflected in the amplitude of the heartbeat-evoked brain potential. Psychophysiology, 41(3), 476–482.

Porges, S. W., & Furman, S. A. (2011). The early development of the autonomic nervous system provides a neural platform for social behavior: A polyvagal perspective. Infant and Child Development, 20(1), 106–118.

Quintana, D. S., Alvares, G. A., & Heathers, J. A. J. (2016). Guidelines for Reporting Articles on Psychiatry and Heart rate variability (GRAPH): recommendations to advance research communication. Translational Psychiatry, 6(5), e803.

Richter, F., García, A. M., Rodriguez Arriagada, N., Yoris, A., Birba, A., Huepe, D., Zimmer, H., Ibáñez, A., & Sedeño, L. (2021). Behavioral and neurophysiological signatures of interoceptive enhancements following vagus nerve stimulation. Human Brain Mapping, 42(5), 1227–1242.

Rigato, S., Rieger, G., & Romei, V. (2016). Multisensory signalling enhances pupil dilation. Scientific Reports, 6(1), 26188.

Rominger, C., Weber, B., Aldrian, A., Berger, L., & Schwerdtfeger, A. R. (2021). Short-term fasting induced changes in HRV are associated with interoceptive accuracy: Evidence from two independent within-subjects studies. Physiology & Behavior, 241(113558), 113558.

Seth, A. K. (2013). Interoceptive inference, emotion, and the embodied self. Trends in Cognitive Sciences, 17(11), 565–573.

Shoji, M., Mehling, W. E., Hautzinger, M., & Herbert, B. M. (2018). Investigating multidimensional Interoceptive Awareness in a Japanese population: Validation of the Japanese MAIA-J. Frontiers in Psychology, 9, 1855.

Sim, Z. L., & Xu, F. (2019). Another look at looking time: Surprise as rational statistical inference. Topics in Cognitive Science, 11(1), 154–163.

Skora, L. I., Livermore, J. J. A., & Roelofs, K. (2022). The functional role of cardiac activity in perception and action. Neuroscience and Biobehavioral Reviews, 137(104655), 104655.

Suga, A., Uraguchi, M., Tange, A., Ishikawa, H., & Ohira, H. (2019). Cardiac interaction between mother and infant: enhancement of heart rate variability. Scientific Reports, 9(1), 20019.

Tünte, M. R., Hoehl, S., Wunderwald, M., Bullinger, J., Boyadziheva, A., Maister, L., Elsner, B., Tsakiris, M., & Kayhan, E. (2025). Respiratory and cardiac interoceptive sensitivity in the first two years of life. ELife, 12. 10.7554/eLife.91579

Vabba, A., Porciello, G., Monti, A., Panasiti, M. S., & Aglioti, S. M. (2023). A longitudinal study of interoception changes in the times of COVID-19: Effects on psychophysiological health and well-being. Heliyon, 9(4), e14951.

van den Heuvel, M. P., Kahn, R. S., Goñi, J., & Sporns, O. (2012). High-cost, high-capacity backbone for global brain communication. Proceedings of the National Academy of Sciences of the United States of America, 109(28), 11372–11377.

van den Heuvel, M. P., Sporns, O., Collin, G., Scheewe, T., Mandl, R. C. W., Cahn, W., Goñi, J., Hulshoff Pol, H. E., & Kahn, R. S. (2013). Abnormal rich club organization and functional brain dynamics in schizophrenia. *JAMA Psychiatry (Chicago*, Ill.), 70(8), 783–792.

Viglione, A., Mazziotti, R., & Pizzorusso, T. (2023). From pupil to the brain: New insights for studying cortical plasticity through pupillometry. Frontiers in Neural Circuits, 17, 1151847.

Villani, V., Tsakiris, M., & Azevedo, R. T. (2019). Transcutaneous vagus nerve stimulation improves interoceptive accuracy. Neuropsychologia, 134(107201), 107201.

Wang, C.-A., Blohm, G., Huang, J., Boehnke, S. E., & Munoz, D. P. (2017). Multisensory integration in orienting behavior: Pupil size, microsaccades, and saccades. Biological Psychology, 129, 36–44.

Weijs, M. L., Daum, M. M., & Lenggenhager, B. (2023). Cardiac interoception in infants: Behavioral and neurophysiological measures in various emotional and self-related contexts. Psychophysiology, 60(12), e14386.

Whitehead, W. E., Drescher, V. M., Heiman, P., & Blackwell, B. (1977). Relation of Heart Rate Control to Heartbeat Perception 1,2 (Vol. 2). https://link.springer.com/content/pdf/10.1007%2FBF00998623.pdf

Yates, A. J., Jones, K. E., Marie, G. V., & Hogben, J. H. (1985). Detection of the heartbeat and events in the cardiac cycle. Psychophysiology, 22(5), 561–567.

Zénon, A. (2019). Eye pupil signals information gain. *Proceedings*. Biological Sciences, 286(1911), 20191593.

