## Supplementary information for "Behavioural and physiological evidence for the development of cardiac-exteroceptive integration during the first year of life"

### Supplementary Materials

Table S1. Mean looking time for each synchronous and asynchronous (fast and slow) trials

| | | looking time (mean ± SD, seconds) | | |
|---|---|---|---|---|
| | | synchronous | asynchronous | |
| | | | fast | slow |
| older | systole | 3.68 ± 1.31 | 4.63 ± 1.82 | 4.59 ± 1.52 |
| | diastole | 4.36 ± 1.82 | 4.24 ± 1.67 | 4.40 ± 1.47 |
| younger | systole | 3.99 ± 1.48 | 3.78 ± 1.67 | 4.00 ± 2.40 |
| | diastole | 5.23 ± 2.45 | 5.13 ± 2.57 | 4.83 ± 2.58 |

Table S2. Mean pupil change for 4-s window for each synchronous and asynchronous (fast and slow) trials

| | | pupil change (mean ± SD, mm) | | |
|---|---|---|---|---|
| | | synchronous | asynchronous | |
| | | | fast | slow |
| older | systole | -0.01 ± 0.08 | -0.03 ± 0.12 | 0.01 ± 0.15 |
| | diastole | -0.07 ± 0.11 | -0.01 ± 0.11 | -0.02 ± 0.10 |
| younger | systole | -0.10 ± 0.10 | -0.03 ± 0.07 | -0.03 ± 0.11 |
| | diastole | -0.11 ± 0.12 | -0.03 ± 0.13 | -0.01 ± 0.11 |

Table S3. Results of the linear mixed model predicting looking-time bias (LBI) from HRV and MAIA

| | β | s.e. | t-value | df | p-value |
|---|---|---|---|---|---|
| intercept | 0.65 | 0.065 | 10.0 | 656 | < .001 |
| **age** | **0.027** | **0.010** | **2.7** | **656** | **0.008** |
| **heart rate** | **0.027** | **0.013** | **2.2** | **656** | **0.031** |
| HRV-HF | –0.027 | 0.014 | –1.9 | 656 | 0.054 |
| **HRV-LF** | **0.042** | **0.015** | **2.9** | **656** | **0.004** |
| **MAIA-noticing** | **–0.011** | **0.0028** | **–4.0** | **656** | **< .001** |
| MAIA-not distracting | –0.0079 | 0.0043 | –1.8 | 656 | 0.069 |
| MAIA-not worrying | 0.0034 | 0.0043 | 0.79 | 656 | 0.43 |
| **MAIA-attention regulation** | **0.014** | **0.0041** | **3.4** | **656** | **< .001** |
| MAIA-emotional awareness | –0.0018 | 0.0026 | –0.69 | 656 | 0.549 |
| **MAIA-self regulation** | **–0.012** | **0.0058** | **–2.0** | **656** | **0.046** |
| MAIA-trusting | –0.0056 | 0.0039 | –1.4 | 656 | 0.15 |
| MAIA-body listening | –0.0046 | 0.0041 | –1.1 | 656 | 0.25 |

Figure S1

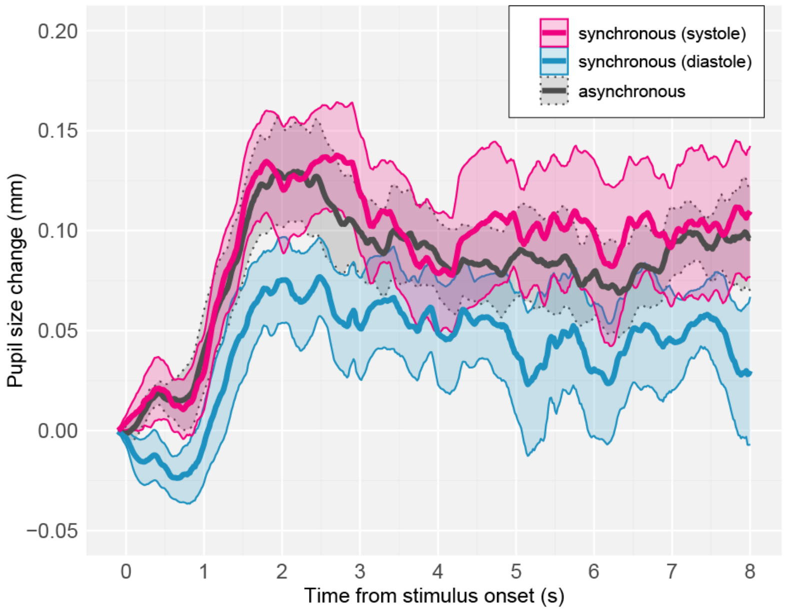


Grand mean time course of baseline-corrected pupil size over the 8 s stimulus presentation in adults. The pink, blue, and grey lines denote systole-locked synchronous (R-peak + 250 ms), diastole-locked synchronous (R-peak + 0 ms), and asynchronous trials, respectively. Shaded bands indicate ± 1 SEM.

#### *Supplementary results and discussion*

We supplementarily examined whether maternal interoceptive sensibility—as measured by eight subscales (noticing, not-distracting, not-worrying, attention-regulation, emotional-awareness, self-regulation, trusting, and body-listening) from the MAIA—predicts infant LBI, under the hypothesis that more interoceptively attuned caregivers scaffold infants' interoceptive integration. We extended the model to HRV by adding these eight MAIA subscale scores. Inclusion of the MAIA variables significantly improved model fit over the HRV-only model ($\chi 2(8) = 31.9$, $p < 0.001$). In this model, in addition to age and LF, attention-regulation emerged as a significant predictor ($\beta = 0.014$, $t(656) = 3.4$, $p < 0.001$), indicating that mothers, who were better able to deliberately focus on their own bodily sensations, had infants with a stronger bias toward asynchronous stimuli. By contrast, higher noticing ($\beta = –0.011$, $t(656) = –4.0$, $p < 0.001$) and self-regulation ($\beta = –0.012$, $t(656) = –2.0$, $p = 0.046$) scores predicted a stronger asynchronous bias (Table S3).

Theoretical frameworks suggest that early interoceptive–exteroceptive integration develops primarily within the caregiving context (Fotopoulou and Tsakiris 2017), and that caregivers' interoceptive sensitivity is assumed to facilitate their inferences about infants' physiological and emotional states and ability to form accurate perceptions of bodily sensations (Montirosso, Mascheroni, and Mariani Wigley 2022). Therefore, we hypothesised that maternal interoceptive sensibility assessed by the MAIA would predict infants' behavioural sensitivity to cardiac–exteroceptive synchrony. However, our findings revealed a more nuanced picture. Specifically, the maternal attention regulation score was positively associated with infants' sensitivity to cardiac–exteroceptive synchrony, whereas the maternal noticing and self-regulation scores were negatively associated. Attention regulation reflects the capacity to focus on internal bodily sensations without being distracted by external stimuli, suggesting a controlled, goal-directed attentional mechanism. In contrast, noticing refers to the automatic and effortless awareness of internal sensations, without involving evaluative or regulatory components. This contrast may be crucial in the caregiving context. Effective caregiving likely requires flexible attention shifting between the caregivers' own internal signals and infants' exteroceptive cues (e.g. crying or facial expressions). Thus, the mothers' ability to regulate their attention, rather than merely notice bodily sensations may support a more adaptive attunement to their infants' internal states. By contrast, heightened noticing may reflect a self-focused attentional style that interferes with social attunement. Supporting this view, Suga et al. (2022) found that in a diaper-changing context, mothers in a control group (who had not received any intervention) showed increases in noticing, that were associated with a greater negative affect, likely due to excessive attention to their own discomfort. Similarly, the self-regulation subscale—which refers to using attention to bodily sensations to manage distress—may also reflect a self-oriented coping strategy, that while potentially beneficial for mothers, could detract from responsiveness to their infants' needs. On the other hand, it is notable that a recent study found positive associations between the noticing, body-listening, and self-regulation subscales of the MAIA and mothers' self-reported engagement in stroking and rocking their infants (Donaghy, Shinskey, and Tsakiris 2024). However, because that study relied on maternal reports of caregiving behaviour rather than objective indices of infants' interoceptive development, its findings are not necessarily inconsistent with the present results. Overall, our findings suggest that not all aspects of interoceptive sensibility support optimal parent–infant coregulation; rather, those related to attentional flexibility may be most conducive to the development of early interoceptive–exteroceptive integration in infancy.

#### *References*


Donaghy, Rosie, Jeanne Shinskey, and Manos Tsakiris. 2024. "Maternal Interoceptive Focus Is Associated with Greater Reported Engagement in Mother-Infant Stroking and Rocking." *PloS One* 19 (6): e0302791.

Fotopoulou, Aikaterini, and Manos Tsakiris. 2017. "Mentalizing Homeostasis: The Social Origins of Interoceptive Inference." *Neuropsychoanalysis* 4145 (June): 1–9.

Montirosso, Rosario, Eleonora Mascheroni, and Isabella Lucia Chiara Mariani Wigley. 2022. "Maternal Embodied Sensitivity: Could Interoception Support the Mother's Ability to Understand Her Infant's Signals?" In *Key Topics in Perinatal Mental Health*, 447–55. Cham: Springer International Publishing.

Suga, Ayami, Yosuke Naruto, Venie Viktoria Rondang Maulina, Maki Uraguchi, Yuka Ozaki, and Hideki Ohira. 2022. "Mothers' Interoceptive Sensibility Mediates Affective Interaction between Mother and Infant." *Scientific Reports* 12 (1): 6273.
